# Predictive Uncertainty in State-Estimation Drives Active Sensing

**DOI:** 10.1101/2023.11.02.565312

**Authors:** Osman Kaan Karagoz, Aysegul Kilic, Emin Yusuf Aydin, Mustafa Mert Ankarali, Ismail Uyanik

**Affiliations:** Electrical and Electronics Engineering, Middle East Technical University, 06800, Ankara, Turkiye; Center for Robotics and AI, Middle East Technical University, 06800, Ankara, Turkiye; Bioengineering Division, Hacettepe University, 06800, Ankara, Turkiye; Electrical and Electronics Engineering, Hacettepe University, 06800, Ankara, Turkiye

**Keywords:** active sensing, weakly electric fish, sensorimotor control, state estimation

## Abstract

Animals use active sensing movements to shape the spatiotemporal characteristics of sensory signals to better perceive their environment under varying conditions. However, the underlying mechanisms governing the generation of active sensing movements are not known. To address this, we investigated the role of active sensing movements in the refuge tracking behavior of *Eigenmannia virescens*, a species of weakly electric fish. These fish track the longitudinal movements of a refuge in which they hide by swimming back and forth in a single linear dimension. During refuge tracking, *Eigenmannia* exhibits stereotyped whole-body oscillations when the quality of the sensory signals degrades. We developed a feedback control model to examine the role of these ancillary movements on the task performance. Here, we show that the proposed model generates fish trajectories that are statistically indistinguishable from the actual fish, implying that active sensing movements are regulated to minimize the predictive uncertainty in state estimation.

## INTRODUCTION

Active sensing is a dynamic process in which animals orchestrate feedback loops to refine their sensory inputs [Nelson and MacIver, 2006, Stamper et al., 2019]. It encompasses a range of intricate movements such as repositioning antennae in arthropods [Dürr and Krause, 2022], eye rotations and saccadic movements in primates [Fuchs, 1967], and exploratory whisking motions of a rodent’s vibrissae [Bush et al., 2016]. Like a skilled technician fine-tuning instruments, these deliberate motor activities prepare an organism’s sensory systems to optimize the perception of specific stimuli types. This active orchestration and selective sensory data acquisition enhance an organism’s understanding of its surroundings, enriching the sensory ‘dialogue’ between the organism and its environment [Schroeder et al., 2010, Stamper et al., 2019]. By meticulously exhibiting active sensing movements, animals obtain diverse and high-resolution information beyond what is achieved via passive reception of the sensory signals in the environment [Biswas et al., 2018, Kö nig and Luksch, 1998, Schuster, 2008]. It is through these active sensing movements that animals achieve stable and precise navigation, engage in complex communication, and effectively utilize their habitat, ultimately enhancing their survival in diverse ecological niches [Jones et al., 2021, Schabacker et al., 2021, Stamper et al., 2019, Jun et al., 2016].

Active sensing is a sophisticated mechanism involving purpose-driven movements. It is not simply a reactionary mechanism but rather an intricate, intelligent interplay between the sensory and motor systems, which seeks to adjust sensory inputs for enhanced perception. Translating this interaction into the “language” of engineering requires building closed-loop models, which encapsulate the processes’ dynamism and responsiveness [Jun et al., 2016, Yeo et al., 2016, Yoon et al., 2018]. Within the proposed modeling framework, active sensing inputs act as disturbance signals, perturbing the motor commands generated by the task controller. This is an intriguing paradox as, despite often being orthogonal and even seemingly contrary to achieving task goals, active sensing movements improve state estimation, which in turn improves task performance.

Active sensing manifests in various forms within the biological systems, each tailored to meet the specific needs of the species employing it. These processes can be categorized into distinct types: active illumination and active sensing through movement [Stamper et al., 2019, Zweifel and Hartmann, 2020]. Active illumination, exemplified in bats, involves the transmission of a signal to the environment and the interpretation of the resulting feedback. By analyzing the timing, direction, and frequency changes in the echoes, bats construct a detailed sensory representation of their surroundings, including the distance, shape, and location of objects [Nelson and MacIver, 2006, Jones et al., 2021, Schabacker et al., 2021]. Meanwhile, the whisker movements in rodents exemplify active sensing via movement. These small mammals continually adjust their whiskers in response to environmental changes, enabling spatial awareness and object detection [Bush et al., 2016, Deutsch et al., 2012]. The diversity of active sensing is further exemplified by cockroaches, which utilize antenna movements to detect obstacles and gather tactile information [Okada and Toh, 2006, Chen et al., 2020], spiders that interpret vibration signals transmitted through their webs for object localization [Nakata, 2010, Blamires et al., 2011], and various animals that dynamically modify their sensory volume through adjustments in physical orientation and focus. By selectively fortifying their sensory inputs, these animals continuously reshape their interactions with the environment [Uyanik et al., 2019, Crimaldi et al., 2022, Claverie et al., 2023, Hofmann et al., 2013].

Weakly electric fish demonstrate a sophisticated approach to active sensing by employing both active illumination and active sensing through movement. Active illumination involves electrosense, where these fish continuously emit an electric field and interpret the resulting feedback signals to construct a representation of their environment. The latter mechanism involves engaging in ancillary body movements, allowing them to modulate the sensory feedback from the environment. They can fine-tune their electric perception through these movements to meet their specific needs. This paper focuses specifically on this second aspect of active sensing in weakly electric fish [Stamper et al., 2019, Uyanik et al., 2019, Biswas et al., 2018, Nelson and MacIver, 2006, Jun et al., 2016, Hofmann et al., 2013].

In this paper, we explore various models of active sensing by movement in animals. One model we investigate is the open-loop model, which characterizes the active sensing signal generator in the central nervous system (CNS) as an open-loop noise source that perturbs the motor commands of the task controller. Another model, recently published in the literature Biswas et al. [2023], is based on the stochastic resonance phenomenon observed in biological systems [Manjarrez et al., 2003, Moss et al., 2004, Ward et al., 2002]. Unlike open-loop models, this model–Biswas et al. (2023)–suggests introducing noise only when the state estimation performance falls below a certain threshold. On the one hand, this model is closed-loop, as the active sensing signals are triggered based on feedback from the state variables. On the other hand, once activated, the output of the active sensing generator is an open-loop noise signal that does not change based on the quality of the state estimation. This model is described as having a mode-switching strategy, where the animal switches between a pure tracking controller and a combined mode, where both the tracking controller and the active sensing generator are active. Our research challenges these traditional models with two distinct approaches. First, we explore the impact of mode switching in a new alternating model, which alternates between a pure tracking and pure active sensing controller based on the state estimation performance. Second, we propose a novel closed-loop active sensing generator model, which continuously injects noise into the sensorimotor control process, inversely proportional to the state-estimation performance.

We examined the active sensing behavior of *Eigenmannia virescens*, a species of weakly electric fish, known for their preference to remain hidden within shelter-like structures in their natural habitats. Specifically, we focused on a unique behavior of these fish, termed refuge tracking, where they track the longitudinal movements of their refuge by swimming forward and backward along a single linear dimension [Uyanik et al., 2019, Yang, 2020, Yared, 2020, Roth et al., 2011, Sutton et al., 2016, Von der Emde, 1999]. To study active sensing in *Eigenmannia*, we analyzed the kinematic responses of the fish during their unconstrained free refuge tracking behavior (see Fig. 1a). Our experimental setup involves a test aquarium with a 3D-printed refuge attached to a linear actuator. A computer-controlled system moves the refuge back and forth within the test setup, implementing predefined stimulus trajectories. The fish’s kinematic responses are recorded using a Near-Infrared (NIR) camera positioned beneath the test setup, capturing detailed information about their tracking response and any active sensing movements. These recorded data are utilized to develop a detailed closed-loop model that accurately captures the intricacies of the refuge tracking behavior and the active sensing mechanisms employed by weakly electric fish.

**Figure 1:**
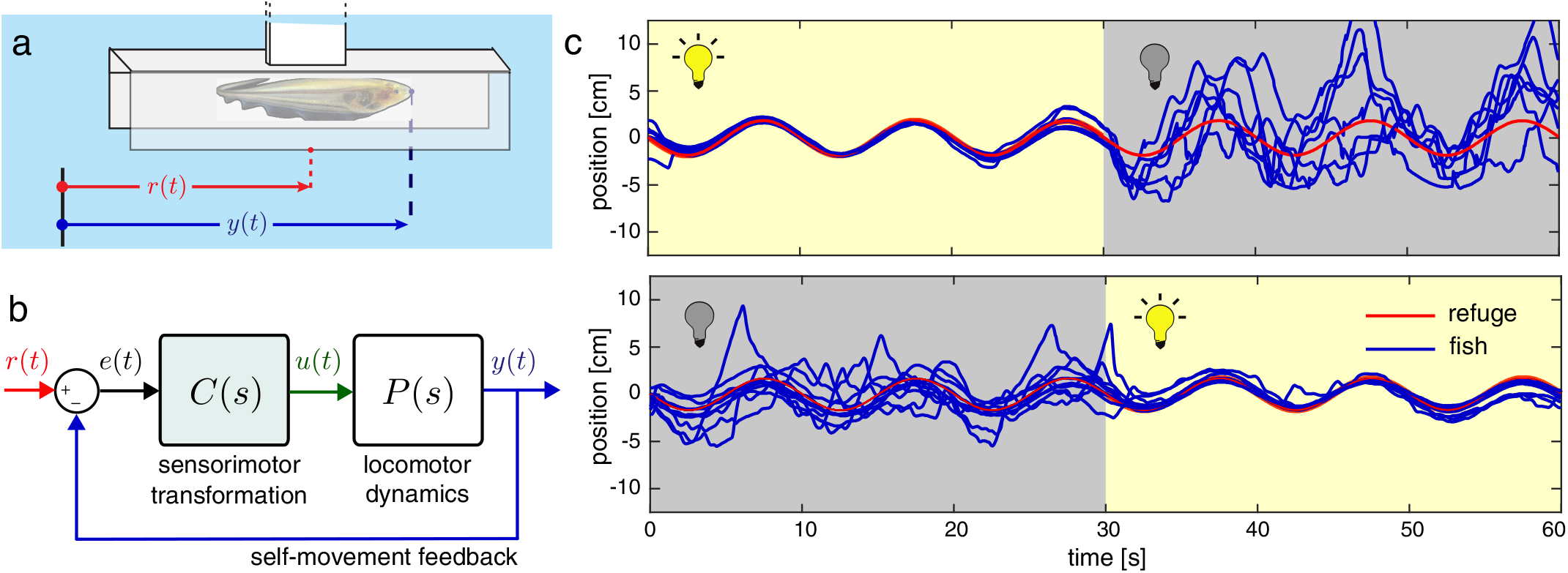
A simple feedback control model for refuge tracking behaviour in *Eigenmannia virescens*. **(a)** *Eigenmannia* tracking a refuge in which it is hiding. **(b)** A simple feedback control model of the refuge tracking behavior. **(c)** *Eigenmannia* exhibits high-frequency movements when swimming in the dark due to increased active sensing movements.(7 trials, light ≈ 300 lux, dark ≈ 0.04 lux)

A vast majority of the literature [Cowan and Fortune, 2007, Roth et al., 2011, Uyanik et al., 2020] encapsulates the sensorimotor transformation–mapping from the sensory input to the motor commands– within a single sensorimotor controller unit (see Fig. 1b). However, these models fail to capture the active sensing movements exhibited by the fish when the quality of the sensory information degrades. Figure 1c illustrates two sample scenarios where the illumination in the environment changed from light to dark or from dark to light. In light, *Eigenmannia* uses both visual and electrosensory systems to detect the relative movement of the refuge. However, in the dark, the fish dominantly relies on electrosensory cues due to the absence of light in the environment. This substantial degradation in the quality of sensory information results in increased active sensing movements that oscillate around the actual tracking movements of the fish, validating the previous results of Stamper et al. [2012].

The key contribution of this paper is the development of a comprehensive mathematical modeling framework that captures the sensorimotor control process employed by the fish during their refuge-tracking behavior. The proposed model encompasses an optimal controller for achieving the behavioral task, an active sensing generator model to drive ancillary body movements, and a state estimator for accurate estimation. This framework provides a versatile platform for exploring different active sensing models and allows for flexibility and adaptability for future research. To ensure the fidelity of our model and enable comparisons between other active sensing generator models, we performed a calibration step using actual fish data obtained during refuge tracking experiments. This calibration procedure helped us to fine-tune the model parameters related to locomotor dynamics and the task controller. This resulted in simulated fish movements that closely aligned with the trial-to-trial variability observed in actual fish behavior. We applied the same approach to assess the predictive performance of various active sensing models for a fair comparison. Our findings demonstrate that the proposed active sensing generator model, which operates based on predictive state-estimation uncertainty, produces active sensing movements that are highly realistic and indistinguishable from the actual fish response unlike the open-loop and stochastic resonance-based models.

## RESULTS

### A feedback control model of active sensing based on predictive uncertainty

We propose that the active sensing behavior in weakly electric fish is intricately linked to the synergistic interactions between the sensory and locomotor systems. To characterize this coupling, we developed a new feedback control model by detailing the simple sensorimotor controller block used in the literature [Cowan and Fortune, 2007, Roth et al., 2011, Uyanik et al., 2020]. Figure 2a illustrates the block-diagram representation of the proposed sensorimotor control model of the fish during refuge tracking.

**Figure 2:**
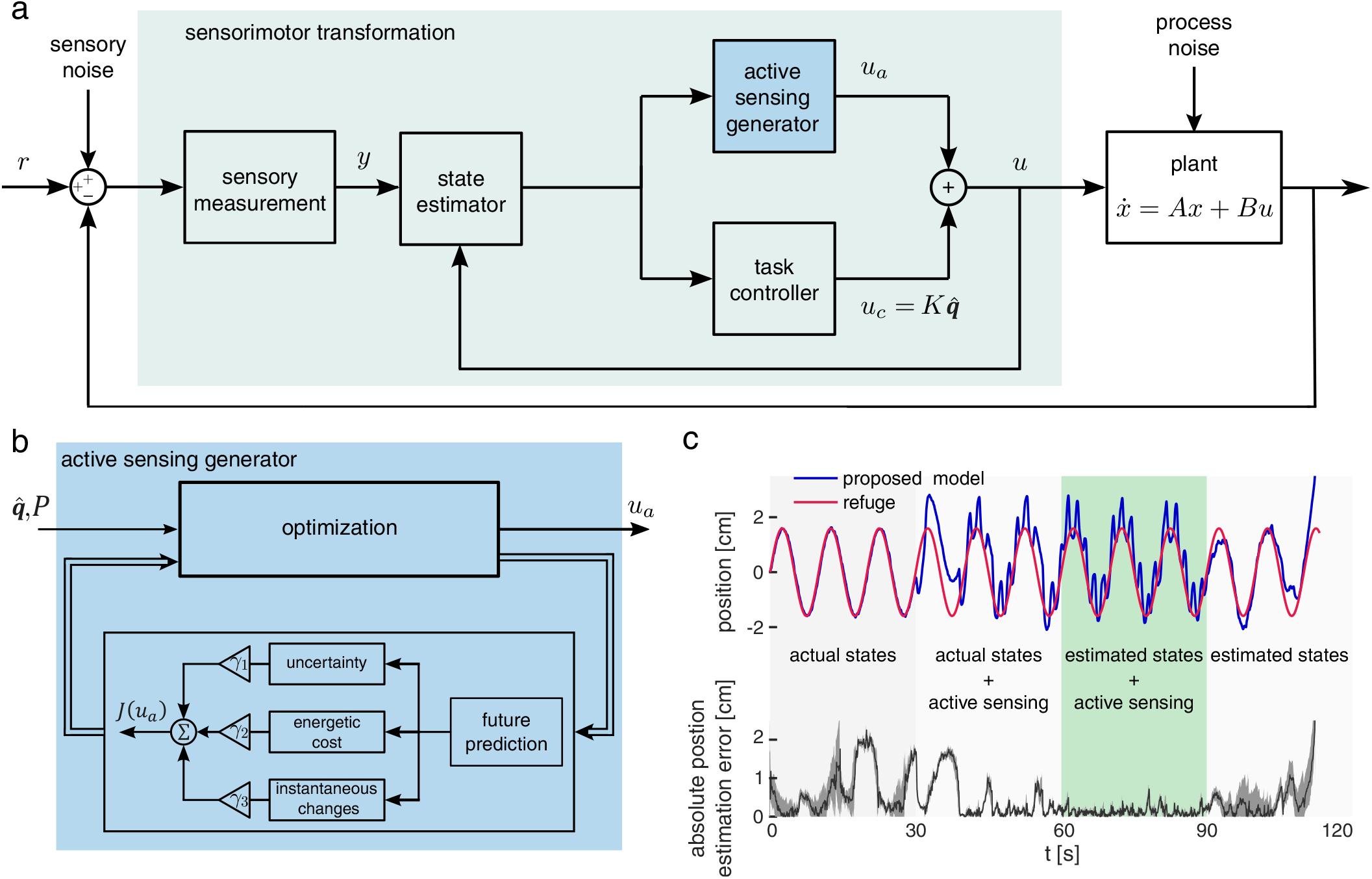
A state-feedback control model of refuge tracking with active sensing. **(a)** The proposed feedback control model of the refuge tracking behavior. The sensory measurement captures the relative movement of the refuge and the fish (contaminated by the sensory noise). The state estimator block estimates the internal states and associated covariance matrix using the efference copy. The task controller and active sensing generator use the state estimates and their covariance matrix to generate the motor commands. The plant, swimming dynamics of the fish, generates movement based on the given motor command. Note that the sensorimotor transformation shown as shaded green corresponds to the sensorimotor transformation of Fig. 1b. We use a more detailed block diagram to analyze the active sensing behavior in this work. **(b)** Block diagram of the proposed active sensing generation method. This block is implemented as an optimization routine to minimize the predictive uncertainty. See Suppoting Information for a possible neural network-based implementation of this optimization routine. **(c)** Demonstrating the need for active sensing. The first epoch uses the actual states to achieve perfect tracking with substantial state estimation error. The second epoch adds active sensing inputs that improve the state estimation performance. However, feeding back the actual state in these two epochs is impossible in real life. The third epoch uses the estimated states and the proposed active sensing generation together, which is the ideal scenario. Here, the state estimation error is minimal, and the model exhibits sufficiently good tracking. In the final epoch, we turn the active sensing off, and the model quickly diverges from the reference as state estimation error increases.

We begin our modeling effort by choosing an appropriate “plant” dynamics corresponding to the swimming dynamics of the fish. In this study, we use the linear, physics-based parametric model developed by Sefati et al. [2013] as the “plant” in the form:

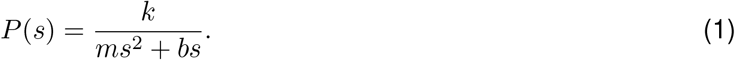

Here, the parameters *m, k*, and *b* denote the mass, gain, and damping coefficients, respectively, while *s* encapsulates the complex frequency in the Laplace domain. In follow-up work, this parametric model was validated during refuge tracking using a data-driven approach [Uyanik et al., 2020].

The plant’s linear nature allows various control theory techniques to be applied. In this study, we implemented a state-feedback control policy for the task controller. We used the pole placement technique to fine-tune the state-feedback gain in the task controller. A critical step in this approach is to calibrate the controller gain to closely match the closed-loop performance demonstrated by the fish in a well-illuminated environment, where the fish needs minimal active sensing. This allows the segregation of the contributions of the task controller and the active sensing generator. The formulation for the state-feedback task controller is as

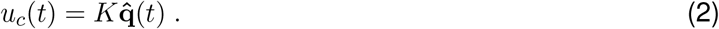

Here, *K* is the state-feedback gain. The vector **q** is defined as **q** = [*p*_*q*_ *v*_*q*_]^*T*^, where *p*_*q*_ and *v*_*q*_ denote the relative position and velocity between the refuge and the fish, respectively. 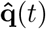 refers to estimated state vector **q**. See Methods and Materials for details.

The task controller, which works based on state feedback, requires an accurate estimation of the system’s internal states, such as the relative movement between the refuge and the fish. The foundational element of the state estimation process is the sensory measurement. To this end, we adopted the electrosensory system model of the weakly electric fish developed by Kunapareddy and Cowan [2018]. This sensory model was also recently used by Biswas et al. [2023] to develop an active sensing generator model for the weakly electric fish. This model aims to capture the electrosensory representation of the external world using a nonlinear measurement function. Mathematically, this sensory model is in the form of

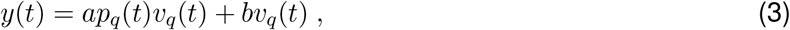

where *a* and *b* are parameters describing the characteristics of the sensory scene around the fish. Note that as the relative velocity, *v*_*q*_ → 0, the information associated with *p*_*q*_ disappears, rendering the system “unobservable”. A detailed mathematical discussion about the observability issue for this sensory model can be found in Kunapareddy and Cowan [2018]. Such a scenario underscores the inevitable need for the use of active sensing strategies, as the sensory scene is unobservable otherwise.

State estimation block plays a crucial role in making the sensory information derived from electroreceptors coherent and usable for the task controller. In other words, the state estimator works as a mapping from the sensory measurements to the neural representation of the sensory information. Due to the inherent nonlinearity of the sensor model, we employed the Extended Kalman Filter (EKF) as our state estimator structure. The EKF is well-suited to generate optimal state estimates given a nonlinearity in the sensor or the process model. By utilizing EKF, we process and interpret the sensory measurements to obtain optimal state estimates and their covariance matrix, which will later be used for the task controller and the active sensing generator as depicted in Fig. 2a.

A key point about the state estimator is that the state estimation accuracy depends critically on the observability of the associated states. One major drawback of the electrosensory model of weakly electric fish (see equation 3) is that the system becomes unobservable when the relative velocity between the refuge and the fish is close to zero, as mentioned before. Intuitionally, this is expected as the electrosensory system of these fish exhibits high-pass filtering characteristics, generating weak or no response when the sensory scene is stationary or at very low speeds. One scenario that causes this problem is when the fish exhibits ideal smooth-pursuit tracking–the velocity of the fish matches that of the refuge. In this case, the state-estimation performance deteriorates, and the fish trajectories deviate from the refuge, as the performance of the state-feedback control depends on the quality of state estimation. The active sensing generator model helps to stabilize this relationship by injecting a purposeful disturbance to the motor commands.

The active sensing generator works in parallel with the linear state-feedback task controller (see the block diagram in Fig. 2a). The goal of the active sensing generator is to perturb the system to recover the observability. Thus, an ad-hoc model for the active sensing generator would be an open-loop noise source injecting noise into the system. However, the previous works clearly showed that animals adopt different stereotyped active sensing actions, whose form and strength vary depending on the quality of the sensory information [Uyanik et al., 2019, Comertler and Uyanik, 2021]. Motivated by the successful results of these works, we model the active sensing generator as a closed-loop regulator, which injects a targeted active sensing signal, trying the minimize the predictive uncertainty in the state estimations. To make it more practical for real-life scenarios, we conceptualized the active sensing generator as an optimization problem considering also the effects of energetic costs and the instantaneous changes in the active sensing input, *u*_*a*_. Consequently, the objective function governing the active sensing generator is in the form of

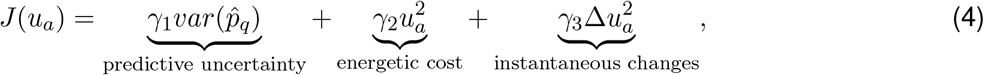

Where 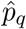 which is a function of *u*_*a*_ is the future prediction of relative position, and *γ*_*i*_’s are the associated weights of the objective function. In our implementations, we used MATLAB’s (MathWorks, Natick, MA) “fminbnd” optimization routine to determine the optimal active sensing input, *u*_*a*_, at each time step. Figure 2c provides a block-diagram representation of the optimization scheme. Detailed information can be found in the Methods and Materials section.

To evaluate the predictive uncertainty—first term in equation 4, we utilize the posterior estimate of the covariance matrix derived by the EKF, which we refer to as the “predictive uncertainty” matrix *P* for brevity. This matrix is instrumental in quantifying the uncertainty, i.e., variance, in the fish’s predictions about its state. Note that a complete state vector associated with *P* may include the global position of the refuge and the fish to perform the simulations. However, some of this information may not be available to the fish. For instance, fish can not infer the global position of the refuge but only its relative position to itself. Also, the unobservability of the system is intrinsically linked to the relative position *p*_*q*_, as mentioned before. Therefore, we only use the variance term associated with *p*_*q*_, the relative motion of the refuge and the fish to evaluate the predictive uncertainty. This approach allows us to accurately capture and incorporate the fish’s sensory limitations into the first cost term of equation 4, thereby enhancing the practicality of our model.

The efficacy of the proposed simulation framework and the active sensing generator model is demonstrated in Fig. 2c. Here, we segmented our simulations into four epochs to show different aspects of the active sensing phenomenon. In the first epoch, we bypassed the state estimator in Fig. 2a and directly fed the actual state estimates to the task controller. Note that this is not possible in a real-life scenario since the controllers can only access an estimate of the states based on sensory measurements. As expected, the fish model performs accurate refuge tracking in this epoch, perfectly matching the reference motion’s frequency, gain, and phase. However, the state estimator fails to infer the states as the system loses observability as the relative velocity *v*_*q*_ approaches zero. Besides, the variance of the state estimation increases substantially. In the second epoch, we keep using the actual states but add an active sensing signal to minimize the predictive uncertainty in state estimations. Therefore, in the second epoch, we observe a drastic decrease in the variance of the state estimation error despite considerable peaks in the magnitude of the state estimation error. This is mainly because active sensing signals directly perturb the ideal tracking input, as the task controller uses actual state information. The third epoch reflects the premise of the current paper, where the task controller uses state estimations, and the active sensing generator seeks to improve the state estimation performance. In this epoch, the magnitude and variance of the state estimation error are both drastically reduced. This, of course, results in additional high-frequency oscillations in the tracking response of the fish. The last epoch shows how the system would respond if the active sensing generator is turned off, while the system uses estimated states to generate control signals. In this scenario, both the magnitude and variance of the state estimation error diverge. As a result, the model loses track of the refuge after a certain amount of time.

### Weakly electric fish use predictive uncertainty to regulate its active sensing movements

The previous section describes the details of our feedback control model that captures the contribution of active sensing in behavioral control. Our results showed that the proposed active sensing generator successfully achieved the task objectives. Despite this success in terms of a design problem, our ultimate goal is not to achieve a perfect active sensing generator that guides the task controller. Instead, we seek to demonstrate if the proposed active sensing generator model and the associated feedback control system can explain the actual fish data. To achieve this, we conducted extensive computational analysis in order to validate the proposed model’s performance in predicting the actual behavior of fish. The simulated feedback control model yields fish trajectories that are statistically indistinguishable from actual fish movements.

To accurately quantify fish behavior, mainly when active sensing movements are prominent, we conducted experiments with *N* = 3 fish under dim light conditions, specifically with illumination around 50 *lux*. During these experiments, we moved the refuge with a constant-frequency sinusoidal input of 0.10 *Hz*, 0.15 *Hz*, or 0.25 *Hz*. Each experiment was repeated 10 times, and we recorded the trajectories of the fish in response to these stimuli. Detailed information about the experimental procedures can be found in Methods and Materials section. The data obtained from these experiments played a critical role in tuning the parameters for our model, ensuring that it closely aligns with the observed behavior of the fish.

We used data obtained from the experiments conducted in well-illuminated conditions for parameter selection in the task-level controller (see Methods and Materials for details). However, our focus shifted to trials conducted under dim light conditions to identify the parameters crucial for the active sensing generator. In these settings, fish exhibit increased active sensing movements. We identified the parameters associated with the sensory scene which are given in equation 3, weights for the cost function given in equation 4, and the parameters related to Extended Kalman Filter (EKF). The values for these parameters were deduced through an optimization routine aimed at minimizing the differences between simulated fish data—derived from stochastic Monte-Carlo simulations with N=1,000 iterations—and the actual observational data of individual fish exposed to specific stimulus frequencies of 0.10 *Hz*, 0.15 *Hz*, or 0.25 *Hz*.

To quantify the difference between our model’s predictions and the actual fish behavior, we employed the Maximum Mean Discrepancy (MMD) method, presented by Gretton et al. [2012]. The MMD offers a reliable way to compare complex distributions without assuming any specific forms regarding the structure of the distributions. An optimization algorithm, detailed in Methods and Materials section, was then utilized to determine the optimal set of parameters that minimizes this distance metric to conduct the parametric system identification. This methodological approach ensures that our model aligns with the observed behaviors of the fish under various conditions.

Figure 3 illustrates the simulation results of the proposed model after fitting the model parameters to the actual fish response. Figure 3a shows the positional data of the fish from 10 different trials and the simulated fish position obtained through our proposed feedback control model. The simulated fish using the estimated model closely replicates the actual fish’s trajectories. Note that the fish has an inherent trial-to-trial variability in its behavioral responses despite using identical stimuli. Our results show that the outputs of the predictive uncertainty model stay well within this trial-to-trial variability.

**Figure 3:**
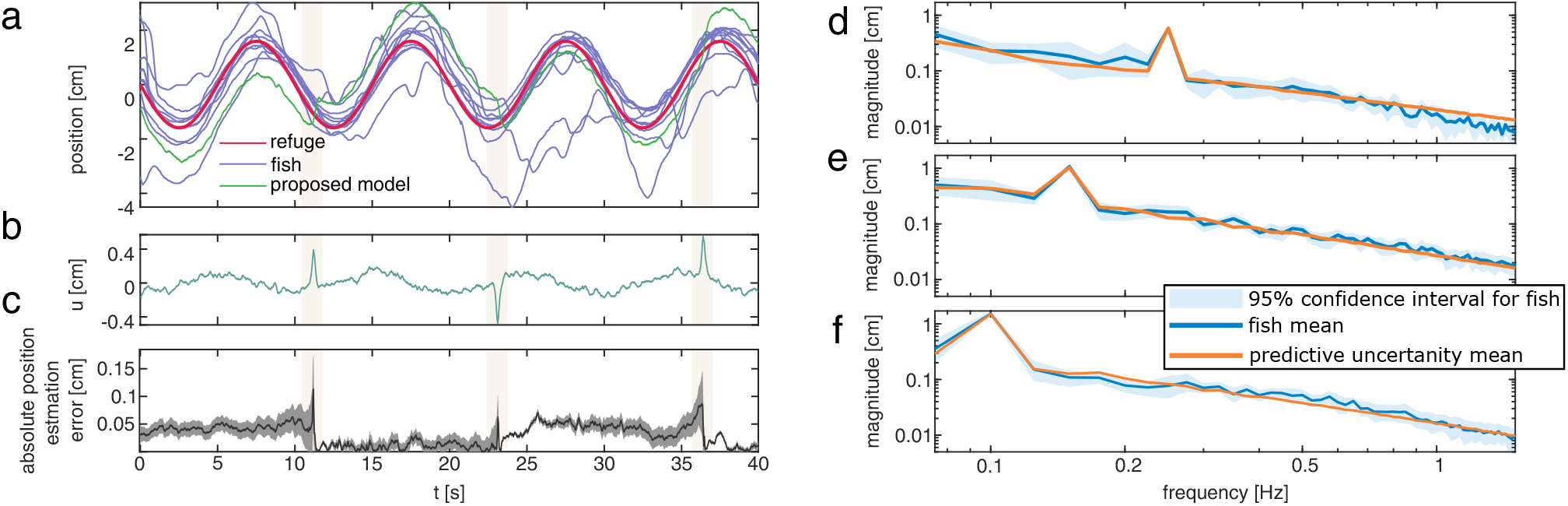
Proposed model vs. fish #2. **(a)** The trajectories (10 replicates) of an actual fish tracking a refuge moving at 0.1 *Hz* and the response of simulated fish using active sensing based on the proposed predictive uncertainty model. **(b)** The result of the total input *u* for the simulated fish in (a). **(c)** State estimation error and associated variance term multiplied by 10 depicting the state estimation quality for the simulated fish in (a). The frequency domain responses of the fish #2 and our model at **(d)** 0.1 *Hz*, **(e)** 0.15 *Hz*, and **(f)** 0.25 *Hz*. The blue-shaded region represents the bootstrap confidence intervals for the mean of the fish response, capturing the %95 range. The solid blue line is the mean response of the fish for the 10 replicate trials for each frequency. The solid orange line represents the mean of our model’s outputs, obtained from 1, 000 random realizations. The response of the proposed model lies within the confidence interval of the actual fish, suggesting that it is indistinguishable from the data of the real fish.

One interesting phenomenon is that the simulated fish generate some fast movements that momentarily deviate the fish from the refuge movements (see shaded areas in Fig. 3). Similar stereotyped movements have also been reported in the literature, yielding a categorical behavior termed as *reset* [Uyanik et al., 2019]. Here, we investigated the underlying reason behind such instantaneous actions preferred by the fish, interestingly also captured by our model. Figure 3b-c depicts the total input *u*, the state estimation error, and the associated variance. Note that these fast movements occur when the active sensing generator undergoes a sudden change. Mathematically, our objective function given in equation 4 aims to prevent such sudden changes in the output of the active sensing generator based on its relative weight in the overall cost. However, the increased variance in the state estimation overcomes the smoothness weight after a certain threshold, yielding another optimal solution that can only be achieved with a sudden change in the control action. Note that the model experiences increased uncertainty in the state estimation error apriori to these sudden control actions.

The time domain plots in Fig. 3 provide an initial understanding about how the proposed model, and hence the active sensing mechanism in fish, works. However, it is quite challenging to quantify the predictive power of the proposed model in terms of capturing the details of the actual fish movements. Besides, the periodic nature of the input-output signals motivates us to perform the quantitative analysis in the frequency domain.

Figure 3 presents a complete frequency domain demonstration of an actual fish’s data and the results of the proposed model. To achieve this comparison, we employ the Discrete Fourier Transform on the time-domain data of the actual fish, constructing a 95% confidence interval that encapsulates the trial-to-trial variability observed in the fish’s response. This interval is derived from the data collected across the 10 experimental trials. We also report the mean response of the simulated fish, obtained from 1, 000 stochastic simulations incorporating variations in model parameters like process and measurement noise. Our results show that for all three test frequencies, the results of the proposed model remain within the confidence interval of the actual fish except for a few exceptional frequencies, at which we do not stimulate the fish. In the following section, we will provide further statistical analysis and compare our approach with other methods in the literature.

### Active sensing is a closed-loop phenomenon

This section aims to conduct a comparative analysis of the predictive performance of the proposed model. One simple explanation for generating active sensing movements is introducing an open-loop noise signal that perturbs the system dynamics, enabling observation of the states for state estimation. Open-loop noise generator models may be sufficient for accurate state estimation and successful tracking. The simulation framework we’ve developed allows the implementation of this model as well, with a minor tweak in the block diagram depicted in Fig. 2a. In the case of an open-loop model, the active sensing generator block does not require any input from the state estimator. Instead, it pumps in a Gaussian white noise signal as a disturbance to the output of the task controller. The strength of the open-loop noise signal is tuned to obtain an optimal fit to the actual fish’s response based on the same identification procedure described in the previous section. Here, we evaluate if the open-loop model can capture the active sensing movements observed in actual fish.

Figure 4 shows the results of our comprehensive analysis with the open-loop active sensing generator model. In Fig. 4a, we report the time-domain trajectories of the actual fish, obtained from 10 replicated trials as well as the response of the simulated feedback control model with the open-loop active sensing generator. Adding an open-loop noise signal to the task controller yields noisy oscillations modulated by the low-frequency tracking signal at the stimulus frequency. Figure 4b shows the corresponding state estimation error and variance, quantifying the state estimation performance. The open-loop model yields considerably less state estimation error than no active sensing scenario presented in the last epoch of Fig. 4c. However, the state estimation error and associated variances are quite higher than the results of the proposed predictive uncertainty method (see Fig. 3). Finally, Fig. 4c shows the frequency response plots of the actual fish with 95% confidence interval and the response of the feedback control model based on an open-loop active sensing generator. The key result here is that the responses due to the open-loop model do not fit well into the trial-to-trial variability observed in real fish despite its successful tracking performance in the time domain. Therefore, the open-loop active sensing generator model’s predictive capabilities are insufficient to capture the details of the actual fish movements.

**Figure 4:**
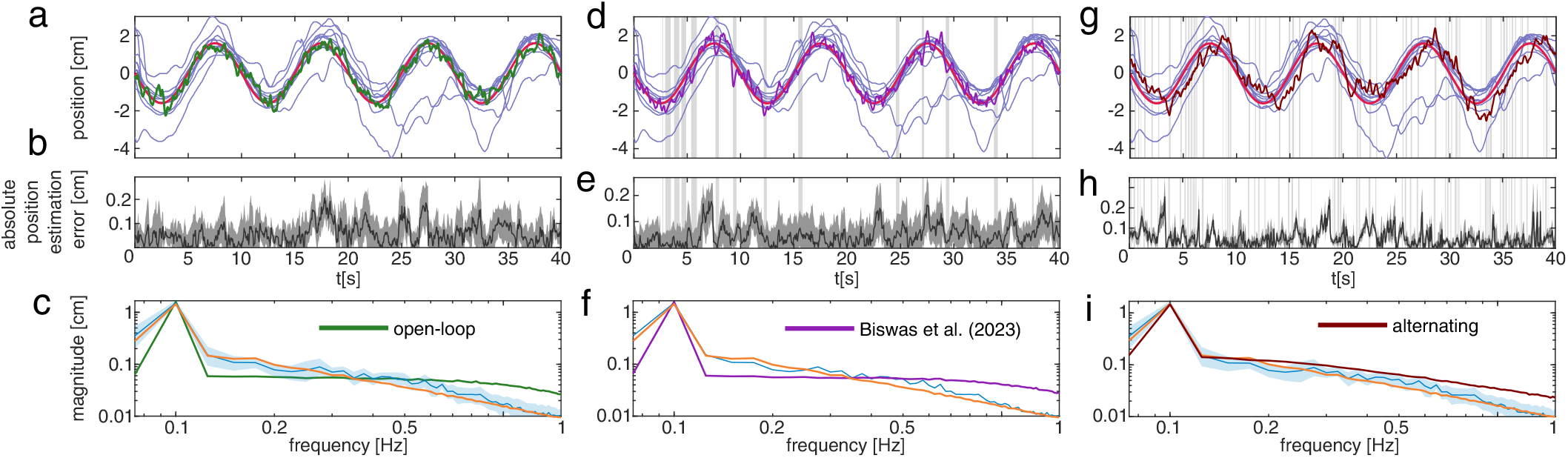
Comparison of open-loop and quasi-open-loop models with fish data. The dark blue and the orange lines represent the mean response fish #2 and the predictive uncertainty model, respectively. The blue regions show the frequency domain response of the fish#2 with 95% confidence interval. On the left, open-loop active sensing model vs. fish #2. **(a)** The trajectories (10 replicates, indicated with the light blue lines) of an actual fish tracking a refuge at 0.1 *Hz* and the response of the open-loop active sensing generator model. **(b)** The state estimation error and the associated variance term multiplied by 10 show the state estimation quality. **(c)** The frequency domain response of the fish with 95% confidence interval and the mean response of the simulation model with an open-loop active sensing generator. On the middle, Biswas et al. (2023) vs. fish #2. **(d)** The trajectories (10 replicates, indicated with the light blue lines) of an actual fish tracking a refuge at 0.1 *Hz* and the response of the active sensing generator model of Biswas et al. (2023). The shaded regions show where the active sensing is off. **(e)** The state estimation error and the associated variance term multiplied by 10 show the state estimation quality. **(f)** The frequency domain response of the fish with 95% confidence interval and the mean response of the simulation model with the active sensing generator model of Biswas et al. (2023). On the right, alternating mode model vs. fish #2. **(g)** The trajectories (10 replicates, indicated with the light blue lines) of an actual fish tracking a refuge at 0.1 *Hz* and the response of the alternating active sensing generator model. The shaded regions show where the active sensing is off and the task controller is on. **(h)** The state estimation error and the associated variance term multiplied by 10 show the state estimation quality. **(i)** The frequency domain response of the fish with 95% confidence interval and the mean response of the simulation model with the alternating active sensing generator model.

Additionally, we compare our findings to a recent model proposed by Biswas et al. (2023), which describes active sensing movements in these fish (see Fig. 4). This approach is primarily based on a different application of the stochastic resonance phenomenon observed in biology and engineering. Unlike open-loop models, Biswas et al. (2023) employ a semi-closed-loop model that activates the active sensing generation signal only when the state estimation performance falls below a certain threshold and deactivates when it surpasses another threshold. In this model, the amount of active sensing input injected into the system is independent of the system’s states. Still, it is triggered based on the state variables, creating a loose dependence on feedback. The simulation framework we’ve developed allows the implementation of this model as well, with minor modifications of the block diagram depicted in Fig. 2a. The application of Biswas et al. (2023) requires the addition of a switch in front of the active sensing generator block, which either enables the injection of an open-loop noise into the control loop or eliminates the contribution of the active sensing generator.

For a fair comparison, we identified the three decision parameters used in Biswas et al. (2023) that specify the active sensing generator, such that the model best fits the actual fish data. Specifically, these parameters are the standard deviation of the Gaussian white noise signal and the upper and lower thresholds on state-estimation performance to switch from active sensing to no active sensing. Note that Biswas et al. (2023) studied active sensing within the context of a stabilization task where the refuge is stationary. In our scenario, we utilize this model within the context of a tracking task where the reference position is dynamically changing.

Figure 4 shows the results of our comprehensive analysis with the active sensing generator model of Biswas et al. (2023). The time-domain data presented in Fig. 4d shows that this model also achieves a sufficiently good tracking performance with extra high-frequency oscillations around the stimulus trajectory. Note that Biswas et al. (2023) differ from the open-loop model in the sense that it can switch off the active sensing generator. The durations where the model switches off the active sensing generator are shaded in gray. Figure 4e shows the corresponding state estimation error and variance, quantifying the state estimation performance. The performance is comparable to the open-loop case as Biswas et al. (2023) approximates the open-loop model as the duration of the shaded regions approaches zero. Figure 4f illustrates the frequency response distribution of the actual fish vs. Biswas et al. (2023). Note that this model also does not generate frequency responses that lie within the trial-to-trial variability of the actual fish.

A fundamental issue with the open-loop model and Biswas et al. (2023) is that they add a Gaussian white noise signal on top of the task controller output. Note that these noise signals appear as a constant line across the frequency axis since they have identical power at each frequency. Therefore, adding such noise profiles can only elevate the magnitude plot of the frequency response of the task controller. This is why we see a nearly flat line beyond the stimulus frequency. This is one of the primary reasons why these models can not capture the frequency response characteristics of the actual fish, which have a decreasing power distribution beyond the stimulus frequency. We anticipate two alternative approaches to remedy the inherent problem with these models. First, one can choose a different strength gain for each active sensing epoch. Specifically, the strength of the noise can vary across time to shape the frequency distribution of the contribution of the active sensing input. This approach adds one more degree of freedom to the active sensing generator to fit the actual fish data. However, this results in more complex models with overfitting issues. Besides, one more degree of freedom will prevent a fair comparison among alternative models. Secondly, we can reshape the overall frequency response by alternating between the active sensing generator and the task controller based on the state estimation performance. The key difference here is that at any time instant, either the tracking controller or the active sensing generator can be active. Our motivation here is that the task controller should not generate a state-feedback-based control signal when the state estimation is poor. It is apparent that the task controller will generate erroneous control actions with poor state estimation. Therefore, this new model alternates between the task controller and the active sensing generator during execution. The active sensing generator is momentarily activated to recover the state estimation and then the task controller keeps regulating the system response.

Figure 4 shows the results of our comprehensive analysis with the alternating controller model extended upon Biswas et al. (2023). Figure 4g shows a successful tracking performance with frequent switches between the task controller and the active sensing generator. As anticipated, when the state estimation performance is poor, the system switches to active sensing and after some time it recovers. This is why we observe frequent active sensing epochs, illustrated with gray-shaded regions in Fig. 4g-h. Note that the model switches to active sensing as soon as the overall state estimation quality drops below a certain threshold. Figure 4i shows the frequency response distribution of the alternating mode model. Despite discrepancies, the alternating mode model exhibits very close performance concerning the trial-to-trial variability of the fish.

We also performed a comparative analysis for the four methods mentioned in this paper across all *N* = 3 fish and the three stimulation frequencies used in the experiments. To ensure an equitable evaluation, we calibrated the model parameters for each scenario to achieve its optimal performance in accurately reflecting the empirical data. We employed a consistent metric to discern active sensing movements, facilitating the quantification of disparities between the frequency response distributions observed in the experiments and those predicted by the models with their optimally adjusted parameters.

Figure 5 illustrates the MMD scores of the four methods across all fish and test frequencies. The proposed predictive uncertainty method yields the minimum MMD score in all fish and almost at all frequencies, suggesting that the frequency distribution is closest to the actual fish data among the four models. The open-loop model and Biswas et al. (2023) yield similar results in all scenarios. It is important to note that the model by Biswas et al. (2023) often operates like an open-loop model, as the active sensing mechanism is frequently engaged (see Fig. 4d). Consequently, the MMD values for the open-loop model and Biswas’s model exhibit high similarity. Meanwhile, the alternating mode model shows performance comparable to, and occasionally rivals, that of the predictive uncertainty method.

**Figure 5:**
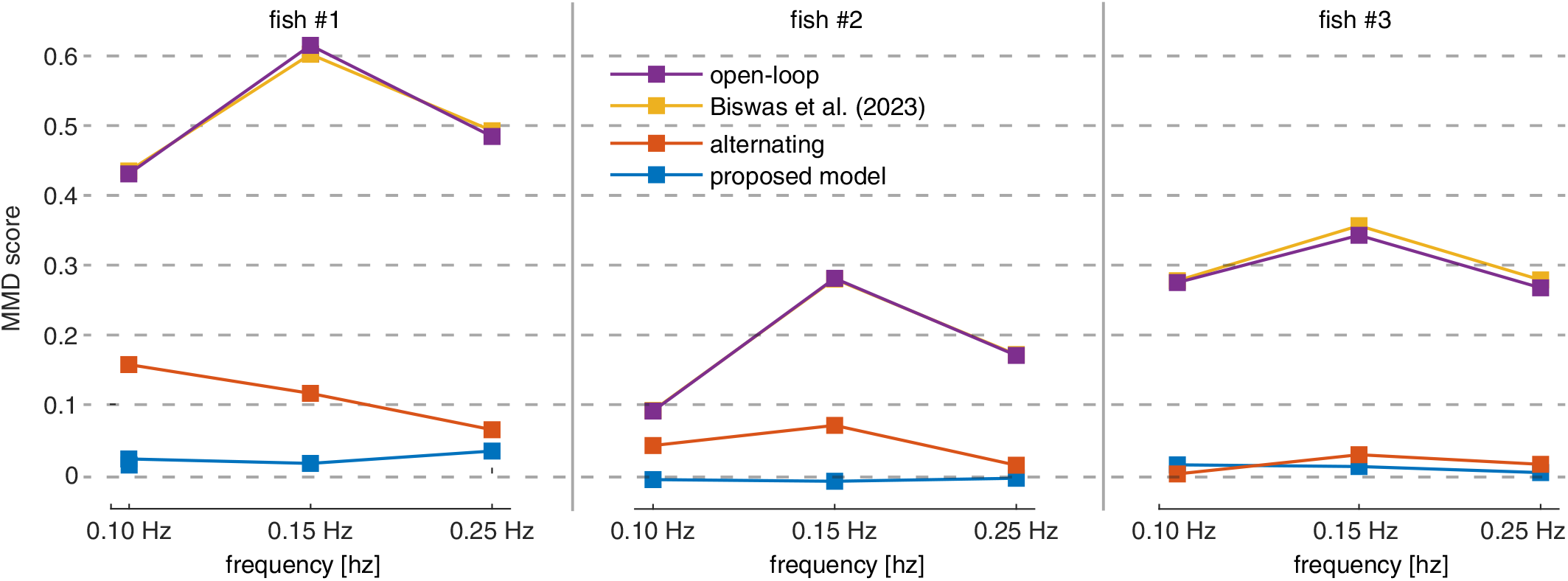
MMD values for the four models for all three fish and the three test frequencies. The proposed predictive uncertainty model exhibits the lowest MMD scores, yielding the closest distributions to the actual fish data. The plots related to fish #2 are given in the main text; see Suppoting Fig. S2. for the results related to fish #1 and fish #3.

The MMD test is instrumental in terms of comparing the distance of distributions. However, we also performed a permutation test, detailed in Methods and Materials to assess if there was a statistical difference between the predicted and actual frequency responses. The null hypothesis suggests no statistical significance between the two distributions. Therefore, a p-value less than 0.05 rejects the null hypothesis, suggesting that the two distributions are statistically different. In the context of system identification, the anticipation is to encounter p-values exceeding 0.05, thereby affirming the lack of statistical significance between the actual and predicted distributions.

Table 1 lists the p-values for all four models for all three fish and the three test frequencies. Biswas et al. (2023) and the open-loop model predictions yield distributions with *p <* 0.0001. This suggests that both of these methods reject the null hypothesis in all scenarios. In other words, one can distinguish the predictions of Biswas et al. (2023) and the open-loop model from the actual trajectories of the fish using statistical analysis despite fitting their parameters to obtain the optimal results–minimum MMD. The alternating mode model, which we proposed as an interim model between Biswas et al. (2023) and predictive uncertainty yields higher p-values, suggesting that it is capable of generating distributions very close to that of the actual fish in most cases. However, the proposed predictive uncertainty model produces frequency response distributions that have no statistical significance as compared to the frequency response distribution of the actual fish in all scenarios. In other words, the predictions of the proposed predictive uncertainty model can not be statistically distinguished from the trajectories of the actual fish.

**Table 1:** Comparison of p-values for all four models and the actual response of the fish.

|  |  | predictive uncertainty | alternating | Biswas et al. (2023) | open-loop |
| --- | --- | --- | --- | --- | --- |
| fish #1 | 0.10 Hz | **0.19\*** | $<0.0001^\dagger$ | $<0.0001^\dagger$ | $<0.0001^\dagger$ |
| | 0.15 Hz | **0.28\*** | 0.01 | $<0.0001^\dagger$ | $<0.0001^\dagger$ |
| | 0.25 Hz | **0.09\*** | 0.01 | $<0.0001^\dagger$ | $<0.0001^\dagger$ |
| fish #2 | 0.10 Hz | **0.86\*** | $<0.0001^\dagger$ | $<0.0001^\dagger$ | $<0.0001^\dagger$ |
| | 0.15 Hz | **0.77\*** | 0.01 | $<0.0001^\dagger$ | $<0.0001^\dagger$ |
| | 0.25 Hz | **0.51\*** | 0.02 | $<0.0001^\dagger$ | $<0.0001^\dagger$ |
| fish #3 | 0.10 Hz | **0.08\*** | **0.35\*** | $<0.0001^\dagger$ | $<0.0001^\dagger$ |
| | 0.15 Hz | **0.13\*** | 0.01 | $<0.0001^\dagger$ | $<0.0001^\dagger$ |
| | 0.25 Hz | **0.32\*** | 0.02 | $<0.0001^\dagger$ | $<0.0001^\dagger$ |

\*Indicates p-values greater than 0.05, suggesting that the evidence against the null hypothesis is not statistically significant at the conventional 0.05 level.
$^\dagger$ Indicates p-values where no permutations were as extreme as the observed data, based on 10,000 permutations.

Note that we used all the data during the parameter estimation process. Therefore, there is a chance of overfitting the data. To avoid this, we conducted cross-validation to reveal if the model performs similarly on test data, which was not used during the parameter identification procedure. We performed cross-validation only on the proposed predictive uncertainty model as the other methods already showed statistical significance despite training them on all data (details in Methods and Materials).

Table 2 summarizes the 3-fold cross-validation results of the predictive uncertainty model. Here, the training and test sets are divided to include nearly equal amounts of data from all frequencies. Note that the proposed model generates distributions with *p >* 0.05 in all test cases, suggesting no statistical significance between the model predictions and the actual fish data, even if the data is entirely novel to the model.

**Table 2:**
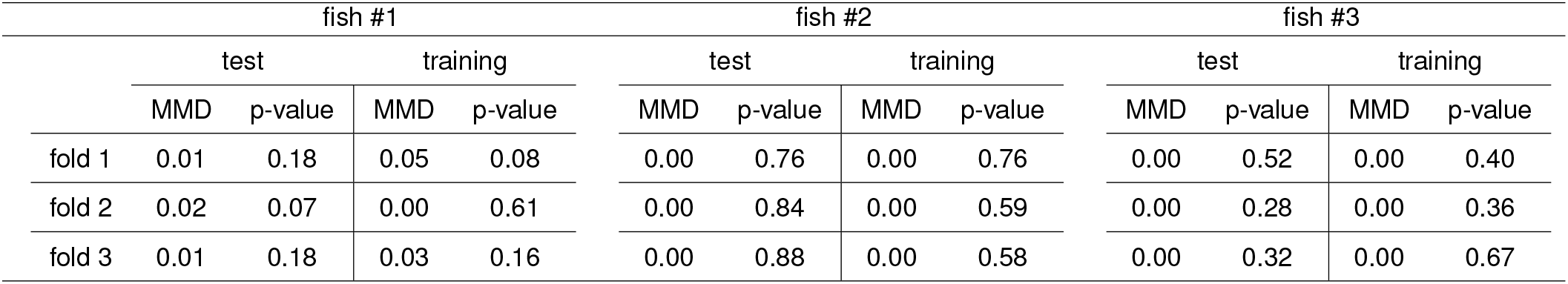
3-fold cross-validation results of the predictive uncertainty model.

## DISCUSSION

Active sensing in animals is a dynamic process in which sensory inputs are delicately interwoven with motor actions, improving behavioral task performance. By initiating ancillary motor actions, animals modify their perception of sensory information, optimizing their understanding of the surrounding environment. However, this active orchestration of sensory-motor coupling that we observe in biological systems challenges our traditional approach to feedback control systems, typically grounded in control theory’s separation principle, which enables engineers to design observers and controllers independently. Embracing and comprehending these violations of separability holds profound implications for advancing our understanding of biological control systems by shedding light on the exceptional adaptability and resilience exhibited by these systems.

Animals demonstrably employ manifold forms of active sensing movements [Bush et al., 2016, Claverie et al., 2023, Crimaldi et al., 2022]. A prevalent hypothesis suggests that active sensing operates as an open-loop system where process noise, or a population of active sensing neurons, introduces broad-spectrum disruptions to the motor commands [Feldman, 2016]. However, evidence of active sensing movements explicitly attuned to the specific behaviors in animals challenges this notion [Yang et al., 2016]. Consider, for example, the minute motoric adjustments made by our fingers to decode the texture of a surface or the distinctive lift-and-shift motion used to deduce an object’s weight; these purposeful actions extend beyond occurrences attributable to random noise interference.

Our study proposes that animals deploy active sensing patterns that are finely tuned to the specific needs of the behavioral task. Specifically, animals generate active sensing movements that minimize the predictive uncertainty of the task-dependent state-estimation performance. To examine this, we studied the active sensing movements generated by weakly electric fish during their refuge tracking behavior. We developed a comprehensive feedback control model for the refuge tracking behavior of the fish, whose parameters meticulously fit the actual fish response through extensive system identification studies, capturing the motion dynamics of the fish. Our results showed that the proposed predictive-uncertainty model generates statistically indistinguishable trajectories for all test cases among the four models we used in our comparative analysis.

The engineering field has extensively explored active sensing, showcasing its effectiveness through numerous working examples [Hinson et al., 2013, Bernard et al., 2020, Hinson and Morgansen, 2013, Cognetti et al., 2018]. However, one may argue that the intricacies of active sensing are not always necessary in engineering. The advent of advanced sensors and sophisticated technologies allows many engineering challenges to be tackled without resorting to the complexities of active sensing. These high-precision and reliable sensors often suffice for fulfilling the required tasks. Conversely, it is important to recognize that natural organisms like weakly electric fish rely on innate mechanisms like active sensing to navigate and interact with their environment. This contrast between engineering and nature emphasizes the adaptability of natural systems and the innovative solutions they have developed over time. While engineering has the advantage of external tools and technologies, nature has evolved to utilize active sensing as an integral part of survival efficiently.

The practical implications of active sensing extend beyond theoretical consequences. It has been observed that individuals with conditions such as autism spectrum disorder, schizophrenia, and attention-deficit/hyperactivity disorder (ADHD) exhibit impaired entrainment to visual or auditory stimuli[Leszczynski and Schroeder, 2019]. These impairments are associated with abnormalities in the neural network responsible for active exploration, involving various cortical and subcortical areas related to specialized functions. Disruptions in this network can result in specific disturbances in active sensing dynamics, such as preparing and executing eye movements and integrating information across saccades. Consequently, these disruptions can serve as diagnostic indicators. For instance, infants who are later diagnosed with autism spectrum disorders tend to focus less on the eyes when viewing faces, while children with dyslexia display alterations in their fixation patterns even before experiencing reading difficulties. Patients with Alzheimer’s disease and mild cognitive impairment exhibit impaired visual search patterns, and individuals with Parkinson’s disease tend to have prolonged fixations when visually exploring complex images. These findings suggest that neurodevelopmental disorders selectively and specifically affect active sensing behavior, indicating early disturbances in the visual and oculomotor systems that alter visual sampling behavior [Schroeder et al., 2010, Leszczynski and Schroeder, 2019].

## METHODS and MATERIALS

### Experimental setup and procedure

We constructed a unique experimental setup (Fig. 6) for this study, resembling a specialized aquarium where the fish engage in tracking behavior within a moving 3D-printed polylactic acid (PLA) refuge. The refuge’s movement is controlled by a high-precision, single-axis linear actuator system powered by a DC motor (Maxon motor 380795, Maxon Group, Switzerland). We used ultra clear glass at the bottom of the aquarium to facilitate the capture of accurate, high-quality, bottom-view images of the fish and the refuge using an NIR camera (Basler ace acA1300-60gm-NIR, Basler AG, Lü beck, Germany) positioned underneath the setup.

**Figure 6:**
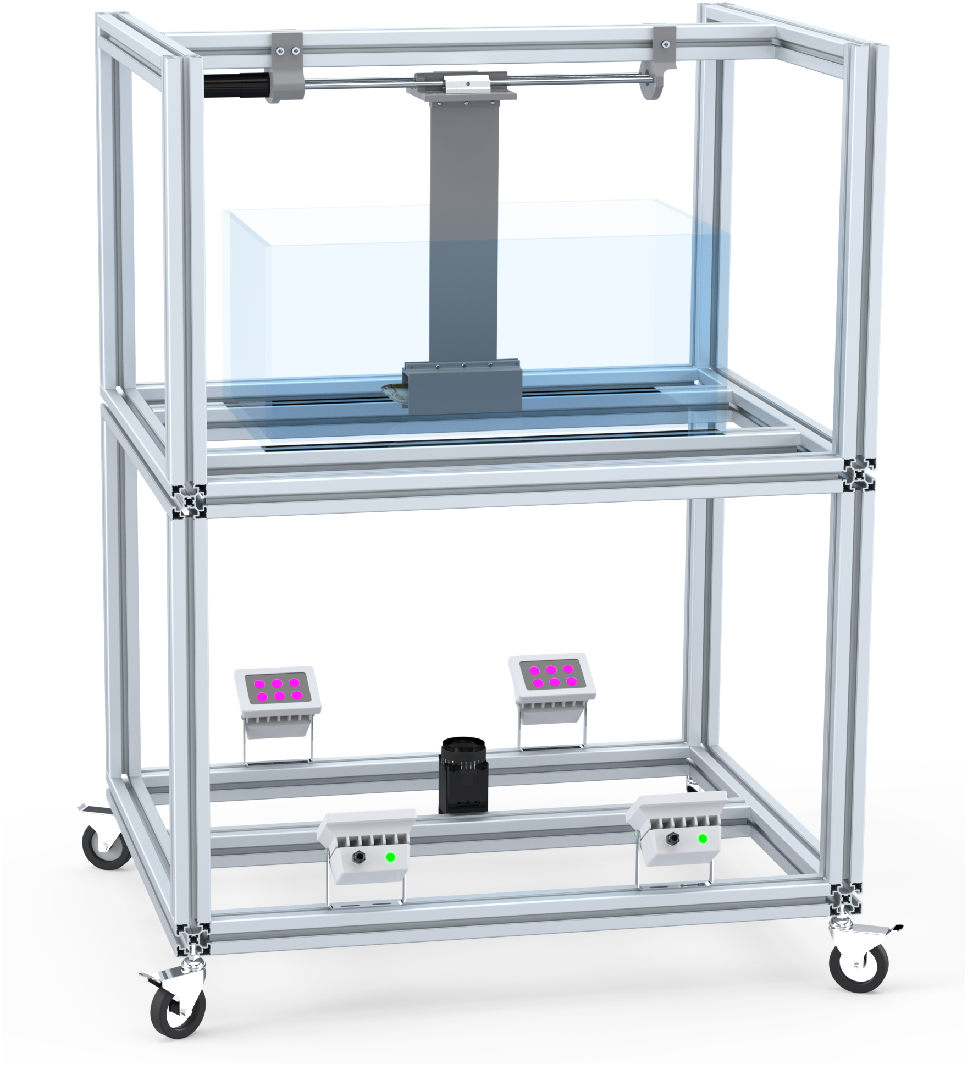
Experimental setup. The fish swims within a refuge attached to a linear actuator on top. The movements of the fish are captured with an NIR camera placed underneath the test area. Infrared LEDs are used to facililatete video recording in the dark.

Our system’s electronic and software architecture runs on the Robot Operating System (ROS). System operations are performed in (nearly) real-time with an online loop frequency of 25 *Hz*. The system’s main software runs on an Nvidia Jetson NX card employing ROS. The Nvidia Jetson board is responsible for executing tasks, including initiating and managing the relevant experiment sequence based on user interface commands, delivering necessary motion commands to the motor driver, and processing feedback about the positions of the motor and the fish. A motor driver (Maxon EPOS4) controls the motion of the linear actuator system based on the commands received from the Nvidia Jetson NX board. The digitized trajectories of the fish are generated by tracking the fish movements from camera records using a custom-built image processing algorithm that works based on template matching.

This study employed three individual weakly electric fish (*Eigenmannia virescens*), aged between 18 and 24 months, to perform refuge tracking experiments. All experiments complied with the guidelines set by the Hacettepe University Animal Experiments Ethics Committee. The water temperature and pH values were stringently maintained throughout the experiments at 25 *±* 1^*°*^*C* and 7.2 *pH*, respectively. The fish were nourished daily with frozen blood worms.

Our research prioritized ethical practices, focusing on minimizing stress on the fish. We did not differentiate between male and female fish used in our trials and ensured no harm was caused to any fish. Before initiating any experiment, we allowed an acclimatization period of 2 to 24 hours, where the fish remained in the experimental setup.

The experiments were performed under light (300 *lux*) and dim light (≈ 50 *lux*) conditions alongside low conductivity (roughly 40 *μS*). Our PLA refuge, measuring 14 *cm* in length and 4 *cm* in width, was utilized for the testing. The refuge was moved with a single constant-frequency sinusoidal input at 0.10 *Hz*, 0.15 *Hz*, or 0.25 *Hz*. These specific frequencies were chosen because they provide a clear separation between active sensing movements and tracking movements, which are not easily distinguishable at higher frequencies due to the relatively high frequency of active sensing movements. The selection of these frequencies was aimed at enabling a thorough validation of the model under study.

Each fish experienced a series of ten repeated trials, each lasting one minute. We used 40-seconds-long steady-state portion of each experimental trial.

## Simulation

### Plant modeling and dynamics

As mentioned earlier, the physics-based parametric model introduced by Sefati et al. [2013] provides a comprehensive framework for characterizing the locomotion behavior of weakly electric fish. Following the established methodology in the field, we adopt the linear musculoskeletal plant in equation 1 for our simulation environment as the “plant” in Fig. 2a.

Uyanik et al. [2020] provide empirical evidence underscoring that while there is inherent and relatively high variation in parameter values within equation 1 among individual fish, there exists a noteworthy consistency in closed-loop performance across different fish. Moreover, they show the resilience of closed-loop performance to mismatches between the plant and controller components by swapping controllers and plants among individuals. This compelling finding suggests that cross-utilization of plant parameters from one fish to another is feasible by establishing a robust feedback mechanism. Building on these insights, our study adopts the parameter values detailed in Uyanik et al. [2020], specifically *m* = 0.0025 *kg, k* = 0.2385 *N/m*, and *b* = 0.0269 *Ns/m*. We use these values as a foundational reference for our analysis.

Describing fish’s state through its position and velocity, represented as ***x*** = [*p*_*x*_ *v*_*x*_] ^*T*^, and making use of equation 1, we can formulate the state-space representation of the plant dynamics as

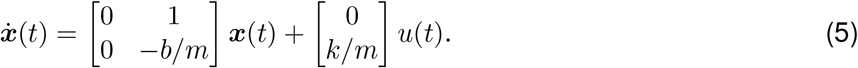

Subsequently, we discretize these continuous equations using a sampling interval of *T* = 0.04 *s* to facilitate discrete-time data acquisition, simulation, and estimator/active sensing formulation. This discrete model that accounts for the intrinsic dynamics of the fish while also incorporating the process noise *ω* is in the form of

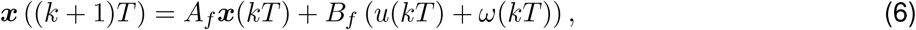

where we assume that process noise is in the form of a zero-mean Gaussian distribution with a standard deviation of *σ*_*w*_, i.e., 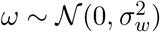.The representation of the refuge, i.e., reference signal, is modeled employing a constant velocity model, a commonly accepted framework in external target tracking applications [Bar-Shalom and Birmi-wal, 1982, Pellegrini et al., 2009, Mahmoudi et al., 2019]. The refuge state, encapsulating its position and velocity, is denoted as ***r*** = [*p*_*r*_ *v*_*r*_]^*T*^. In this paradigm, the dynamic evolution of the refuge system is characterized by:

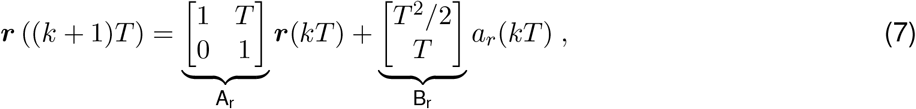

where *a*_*r*_ denotes the acceleration of the refuge.

The proposed combined model augments the existing fish state space by introducing two new variables. This expansion results in the fish’s state space, encompassing its position and velocity attributes, transitioning into a four-dimensional domain. Thus, the state space takes the form of

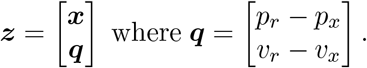

By combining the models given in equation 6 and equation 7, the new state space transition equation can be expressed as

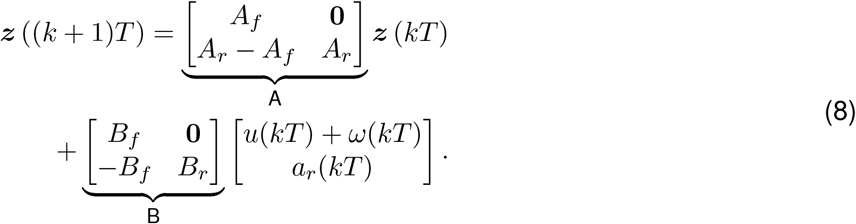

This equation is the system equation used in the simulation environment. Note that *u* is composed of the task-level controller, *u*_*c*_, and the active sensing generation, *u*_*a*_, as *u* = *u*_*c*_ + *u*_*a*_. The schematic representation of this model is depicted in Fig. 2a.

### Controller design and implementation

As previously presented, locomotion behavior relies on two distinct objectives: refuge tracking and enhancement of sensory information. To achieve these objectives, our model assumes that the neural controller comprises two control inputs, a tracking controller and an active sensing generator. Our primary objective is to find models for these controllers that can well explain experimental data. To facilitate this pursuit, we initially undertake the system identification of the task-level controller governing fish behavior within an illuminated environment. This choice is motivated by observing a diminished incidence of active sensing movements in such conditions (see Suppoting Movie S1 and Movie S2). In that respect, we assume that the active sensing generator is either non-existent or active sensing outputs are quite small compared to the tracking controller outputs, thereby allowing us to identify parameters not associated with the active sensing.

To this end, we first concentrate on the whole input-output (from the refuge’s position to fish’s position) dynamics and a second-order model, expressed as

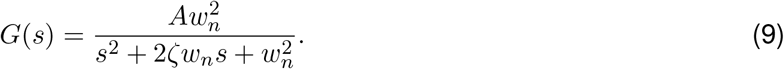

This model, initially proposed by Cowan and Fortune [2007], can effectively capture the input-output dynamics exhibited by the fish. Consistent with Uyanik et al. [2020], we undertake a parametric system identification process to reveal the underlying closed-loop eigenvalues of the dynamics for *N* = 3 fish.

These experiments conducted tests under a well-lit environment, specifically at 300 *lux*. The refuge was subjected to a constant-frequency sinusoidal input, with frequencies set at 0.10 *Hz*, 0.15 *Hz*, and 0.25 *Hz*. For each frequency, a series of 10 experiments was performed, resulting in a total of 30 distinct trials. To facilitate the system identification process, we employed MATLAB’s “greyest” function. Based on all 30 experiments, this method was applied to determine the parameters specified in equation 9 for each fish. The outcomes of this identification process are tabulated in Table 3.

**Table 3:** Parametric identification of the task-level controller. The parameters of a second-order input-output model, in equation 9, were obtained for each fish. We then computed the corresponding eigenvalues, *λ*_1,2_, which are the closed-loop eigenvalues of the system’s transfer function *G*(*s*). This computation was performed after discretizing the system with a sampling time of *T* = 0.04 *s*.

|  | fish 1 | fish 2 | fish 3 |
| --- | --- | --- | --- |
| $A$ | 1.01 | 0.98 | 0.88 |
| $\zeta$ | 0.58 | 0.80 | 1.15 |
| $w_n$ | 9.11 | 7.73 | 9.37 |
| $\lambda_{1,2}$ | $0.77 + 0.24i$ | $0.79 + 0.15i$ | 0.81 |
| | $0.77 - 0.24i$ | $0.79 - 0.15i$ | 0.52 |

Thereupon, we formulated the state feedback mechanism in equation 2 using the pole placement technique for the task-level controller. This approach is combined with the earlier mentioned plant model presented in equation 1, with the objective of aligning the closed-loop eigenvalues to the positions acquired from the parametric system identification process. Assuming *u*_*a*_ = 0 and the state estimator is perfect, then equation 9 stands for the input-output relation of Fig. 2a and we can find *K* in equation 2 using pole placement technique.

### State estimation framework

The equation 8 captures the discrete state transition dynamics of the augmented system, whereas equation 3 details the continuous measurement model. In parallel with our mathematical and data analysis framework, we also discretize this measurement model as

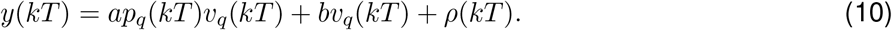

Here, we denote the measurement noise by *ρ*, which follows a zero-mean Gaussian distribution with a standard deviation of *σ*_*ρ*_. Although the system dynamics are linear, the measurement model is nonlinear. To address this nonlinearity in the state estimation approach, we adopt Extended Kalman Filter. The EKF uses equation 8 as the state transition model and equation 10 as the observation model. It is important to note that the fish’s perception of its refuge is modeled with a constant velocity approach, and the inclusion of *a*_*r*_(*kT*) in equation 8 accounts for the uncertainty inherent in the fish’s perception. Thus, it is modeled as a zero-mean Gaussian distribution with a standard deviation of *σ*_*a*_ in the state estimation framework.

### Active sensing generation block

Active sensing extends beyond biological systems, finding significant relevance and application in robotic systems. Notable studies in this area have focused on actively selecting inputs in a closed-loop fashion to maximize observability [Hinson et al., 2013, Bernard et al., 2020, Hinson and Morgansen, 2013]. In our research, we approach the generation of active sensing as an optimization problem, akin to the methodology presented by Cognetti et al. [2018]. The primary goal is to formulate an objective function that concentrates on the future uncertainty of the system in question.

The objective function we propose in equation 4 comprises three components. The initial term represents predictive uncertainty. We quantify this using the posterior estimate of the covariance matrix, *P* ((*k* + 1)*T*) derived by the EKF framework similar to Cognetti et al. [2018]. Salaris et al. [2019] demonstrate that the minimization of a particular norm of *P* ((*k* + 1)*T*) effectively corresponds to the maximization of a similar norm of the constructibility Gramian. This principle suggests a state trajectory can be achieved, culminating in a state estimation with minimal uncertainty. Consequently, an active sensing strategy that aims to minimize *P* ((*k* + 1)*T*) can significantly enhance the state estimation performance of the system under consideration, aligning with the principles of optimal state estimation. At time *kT*, we evaluate *P* ((*k* + 1)*T*) as

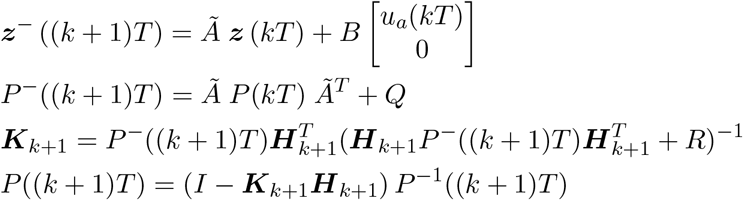

Where 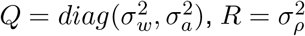, and 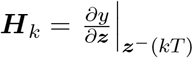.

In this equation set, we consider the new system matrix after applying state feedback to comply with the block diagram in Fig. 2a. Namely, the transformation matrix *Ã* is evaluated as *Ã* = *A* + *BK*. Moreover, as stated previously, the fish’s perception of its refuge is modeled with a constant velocity approach, so *a*_*r*_(*kT*) = 0. It is important to note that we only consider the element of *P* ((*k* + 1)*T*) that corresponds to the uncertainty of *p*_*q*_, specifically the element at the 3rd column and 3rd row.

The remaining terms in equation 4 address eliminating unfeasible solutions for *u*_*a*_(*kT*). The second term, *u*_*a*_(*kT*)^2^, penalizes excessive active sensing input, whereas the final term, (*u*_*a*_(*kT*) − *u*_*a*_((*k* − 1)*T*))^2^, targets rapid changes in the active sensing input. MATLAB’s “fminbnd” optimization algorithm is utilized at each time step to find the active sensing input *u*_*a*_(*kT*) that minimizes this objective function.

### System identification of active sensing

As discussed previously, we identified the parameters non-associated with the active sensing generator and the state estimator based on experimental data collected under well-lit conditions. Here, we aim to identify parameters associated explicitly with the estimation and active sensing operators.

We conducted this system identification process by developing an optimization framework that utilizes MATLAB’s particle swarm optimization algorithm. The optimization’s objective function was constructed around the MMD metric, enabling a comparative analysis between the frequency domain representations of the simulated fish data—generated from stochastic Monte-Carslo simulations (*N* =1,000)—and the actual fish observational data. These observations were specific to individual fish subjected to selected stimulus frequencies, namely 0.10 *Hz*, 0.15 *Hz*, or 0.25 *Hz*. To ensure the stability of the closed-loop system, a penalty function method was integrated into the objective function, a common approach in system identification. The process involved transforming the time domain data of both the simulated and observed fish into the frequency domain using the Discrete Fourier Transform, followed by the evaluation of MMD between these transformed datasets. The optimization algorithm’s goal was to minimize the MMD, thereby aligning the simulated data with the empirical observations as closely as possible. Further details on the implementation of MMD are provided in Methods and Materials.

In the Monte-Carlo simulations, first, we set the task-level controller parameters for the given fish based on the findings presented in Table 3. The initial conditions for the simulated fish—position and velocity—were uniformly distributed around the refuge’s initial state to mirror the natural starting conditions. To eliminate the influence of transient dynamics on the frequency domain data, our analysis concentrated on the final 40 seconds of each 60-second simulation run, a mandatory process in frequency domain system identification and analysis.

The parameters subject to optimization are the scene parameters described in equation 3 and the standard deviations of the noise parameters, specifically *σ*_*ρ*_ and *σ*_*a*_. In our simulations, we kept the standard deviation of process noise constant at *σ*_*w*_ = 0.01. In addition to these, we also adjusted parameters related to the active sensing generation block, which vary based on the specific model in use. For the alternating model and the model proposed by Biswas et al. [2023], we considered three parameters: the threshold pair and the Gaussian input standard deviation. The open-loop model utilized a single parameter, the Gaussian input standard deviation, while the predictive uncertainty model incorporated the *γ*_*i*_’s in equation 4.

## Quantification and statistical analysis

### Bootstrap confidence interval analysis

We implemented a bootstrap resampling technique to assess the reliability of frequency magnitudes derived from the Discrete Fourier Transform applied to the positional data of fish in our experiments. This method entailed generating 10, 000 bootstrap samples from the frequency data, with each sample being drawn with replacement to preserve the size of the original dataset. We computed the mean magnitude across all frequencies for each resampled dataset to form a distribution of mean values. The 95% confidence intervals for the mean magnitudes were then established using the 2.5*th* and 97.5*th* percentiles of this bootstrap distribution. This technique allowed us to infer the trial-to-trial variability and confidence of the frequency components detected in the positional data without relying on parametric assumptions, which is particularly advantageous for non-Gaussian or small sample size data.

### Maximum mean discrepancy evaluation

The Maximum Mean Discrepancy (MMD) is a non-parametric statistical metric that quantifies the difference between two probability distributions [Gretton et al., 2012]. This metric finds extensive application in machine learning, particularly in kernel methods and generative adversarial networks (GANs) training. Additionally, its utilization is notable in the field of bioinformatics [Hao et al., 2024, Ghosh et al., 2023, Wang et al., 2019]. Borgwardt et al. [2006] were pioneers in demonstrating the efficacy of MMD-based two-sample tests across three key data integration tasks: evaluating cross-platform comparability of microarray data, facilitating cancer diagnosis, and implementing data-content-based schema matching for distinct protein function classification schemas. Their studies, including those with high-dimensional data, revealed that MMD excels in identifying samples originating from the same distribution, surpassing traditional methods such as the t-test, multivariate Kolmogorov-Smirnov test, multivariate Wald-Wolfowitz test, and the Biau & Gyö rfi test. Furthermore, they proposed the application of MMD in validating computational simulations of biological processes—a methodology we employ in this study. Herein, MMD serves as a robust metric for comparing simulated fish data against experimental observations, distinguishing itself by accommodating complex distributions without necessitating specific form assumptions, thereby enhancing the assessment of model accuracy and fit.

The MMD is a measure of the distance between two probability distributions, ℙ and ℚ, based on the mean embeddings in a Reproducing Kernel Hilbert Space (RKHS). The RKHS is a space of functions endowed with a kernel function *k*, which is symmetric and positive definite. This kernel function implicitly maps data to a high-dimensional feature space and measures the similarity between data points. The kernel embedding of a distribution ℙ is a function *μ*_ℙ_ in the RKHS defined by the expectation of the feature map associated with *k, μ*_ℙ_ = 𝔼_*x*∼ℙ_[*k*(*·, x*)]. This embedding captures all the moments of the distribution, assuming the kernel *k* is characteristic [Park et al., 2016, Sriperumbudur et al., 2008]. For two distributions ℙ and ℚ, their MMD is defined as the RKHS norm of the difference between their embeddings MMD(ℙ, ℚ) = ∥*μ*_ℙ_ − *μ*_ℚ_∥_ℋ_.

In practice, we don’t have access to the distributions directly but rather samples from them. Given Samples 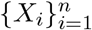 from ℙ and 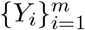 from ℚ, the empirical estimate of the squared MMD is

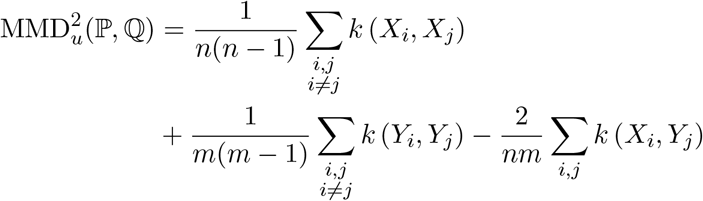

This expression is computationally feasible and does not require explicit computation of the feature maps. It involves only the kernel function evaluations, which are often much simpler to compute. For the results presented in the study, we used this formula and the Gaussian kernel, which is a characteristic kernel, 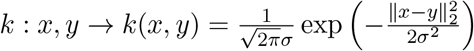. We set the kernel bandwidth as *σ* = 1. Note that since this is an unbiased empirical estimation of MMD, it can take negative values if the true MMD value is close to zero. Those negative values are replaced by zeros while presenting the results.

### Permutation-based p-value calculation for two-sample test

To determine the statistical significance of the MMD between two samples, we employed a permutation test. This non-parametric approach involves combining the samples and randomly shuffling the pooled observations to generate new sample pairs. For each of the *B* permutations, the MMD statistic is recalculated, creating a null distribution of MMD values under the hypothesis that both samples originate from the same distribution. The p-value is then estimated as the proportion of permuted MMD statistics that exceed the observed MMD, with a low p-value indicating a significant difference between the distributions of the two samples. In our study, we set ℬ to 10,000 permutations to ensure an accurate estimation of the p-value. This approach is particularly advantageous in biological data analysis, where the underlying distributions may not conform to standard parametric assumptions, thus necessitating a flexible and assumption-free test.

### Cross-validation for model performance assessment

To evaluate the predictive performance of our model and illustrate that it does not suffer from overfitting in the parameter estimation step, we implemented a cross-validation strategy. Cross-validation is a robust statistical method used to assess the generalizability of a model to an independent dataset. Specifically, we employed a 3-fold cross-validation approach for the proposed predictive uncertainty model.

The dataset was partitioned into three distinct folds, ensuring each fold contained a balanced representation across all stimulus frequencies. This is crucial to maintain the integrity of the cross-validation process, as it prevents the model from being tested on unrepresentative subsets of data.

For each cross-validation fold, the model was trained on two-thirds of the data (the training set) and tested on the remaining one-third (the test set). The training phase involved adjusting the model parameters to minimize the distance between the model outputs and the observed data as described previously. The test phase then evaluated the model’s performance on the test set, which was not exposed to the model during the training phase.

The model’s performance was quantified using the MMD metric, comparing the distribution of the model’s predictions to the actual fish data. As described in the previous section, a p-value was computed to assess the statistical significance of the discrepancy between these distributions.

## ACKNOWLEDGMENTS

This work was supported by The Scientific and Technological Research Council of Turkiye (TUBITAK) under Grant No. 120E198 awarded to Dr. Ismail Uyanik. The authors also thank eLife for the Ben Barres Spotlight Award 2020 awarded to Dr. Ismail Uyanik.

